# Regulation of spikelet number during wheat spike development

**DOI:** 10.64898/2026.01.07.698073

**Authors:** Chengxia Li, Kun Li, Chaozhong Zhang, Jorge Dubcovsky

**Affiliations:** University of California, Davis, CA 95616, USA; Howard Hughes Medical Institute, Chevy Chase, MD 20815, USA

## Abstract

Wheat produces unbranched inflorescences (spikes) composed of smaller inflorescences (spikelets) as their fundamental building units. The spikelet number per spike (SNS) is a major determinant of grain yield and the gene networks that regulate this trait are the focus of this review. Spikelet development starts with the transition of the shoot apical meristem into an inflorescence meristem (IM) that produces lateral spikelet meristems (SMs). The rate at which SMs are produced and the timing of the IM transition into a terminal spikelet (IM→TS) determine the final SNS. These two traits are regulated by genes expressed in the IM (e.g. meristem identity genes), as well as by the amount of FLOWERING LOCUS T1 (florigen) transported from leaves to developing spikes. Spikelet number can also be increased by the production of spikes with supernumerary spikelets (SS) or branch-like structures that resemble small spikes. Mutations that promote a reversion from SM to IM identity can induce the formation of SS or branches. Initial efforts to incorporate these mutations into commercial wheat varieties have faced trade-offs in fertility and grain weight, which will require additional research and breeding efforts. Meanwhile, genes and allele combinations that increase SNS without affecting the number of spikelets per node have been identified and are being deployed in wheat breeding programs. Recent spatial transcriptomics, single-cell analyses, and multi-omics studies of wheat spike development are accelerating the discovery of new genes affecting SNS and enhancing our ability to engineer more productive wheat spikes.

## Introduction

Inflorescences bear flowers and fruits, and their architectures affect yield in crop species. In the grasses, the basic reproductive units are small inflorescences called spikelets, which in wheat contain multiple florets (Fig. 1a-c). As a result, grasses have compound inflorescences or synflorescences [1]. To avoid using a specialized nomenclature for the grass inflorescences, we employed an expanded classification system based on the positions of self-contained developmental reproductive modules within the inflorescence scaffold [2]. Both spikelets and flowers are self-contained developmental programs that can be expressed at different positions within the inflorescence and are initiated by MADS-box genes of the SQUAMOSA clade [3,4]. Using this expanded classification system, an inflorescence bearing flowers or spikelets directly attached to the main rachis is classified as a spike (Fig. 1a).

**Fig. 1.**
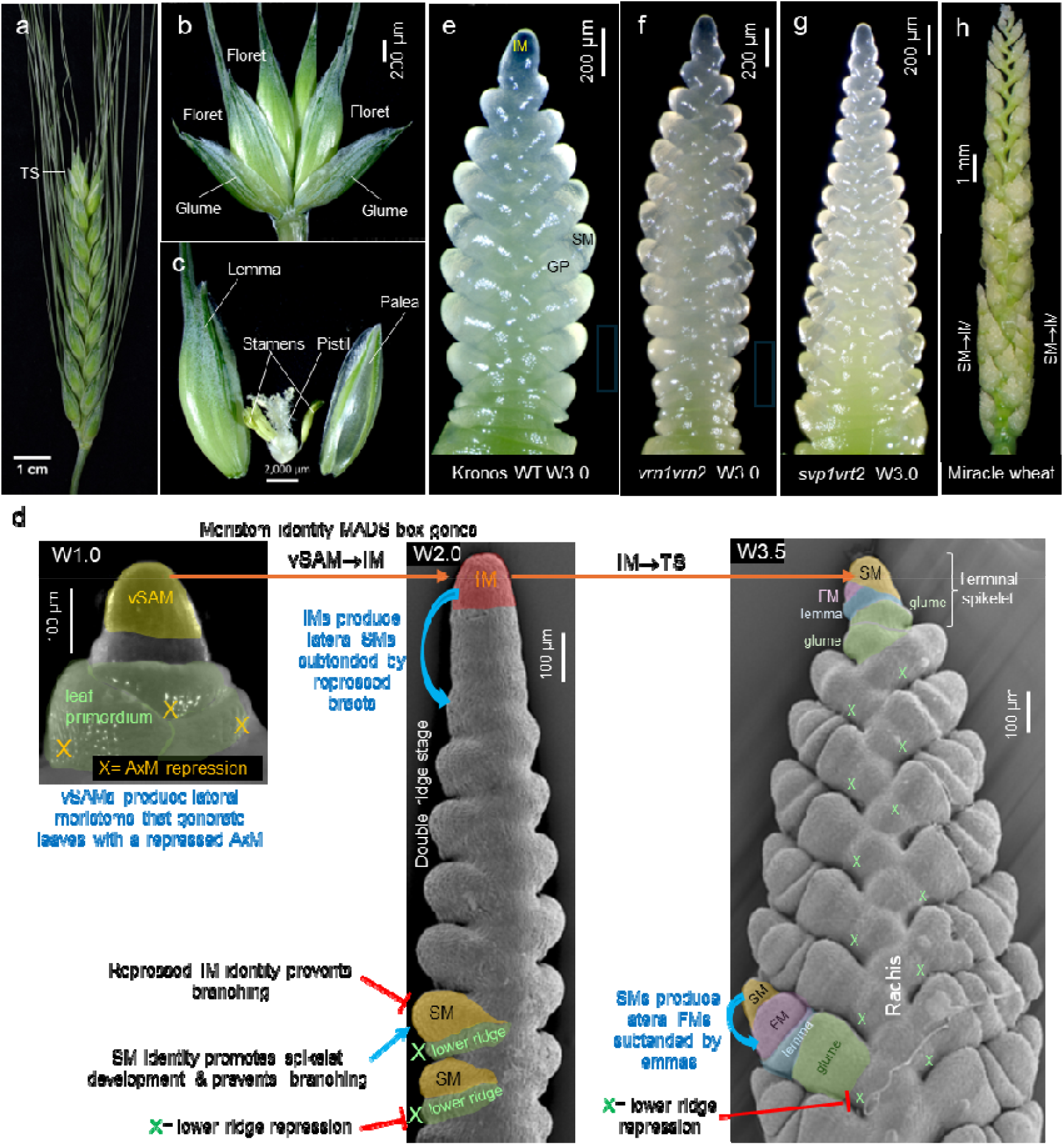
**(a)** Wheat spike. **(b)** Wheat spikelet showing two basal sterile bracts (glumes) and multiple florets. **(c)** Each floret includes a bract (lemma) subtending the floral organs: the palea, two lodicules, three stamens, and a pistil. **(d)** Transitions between different meristem types that shape the wheat spike: vSAM= vegetative meristem, AxM= axillary meristem, IM= inflorescence meristem, SM= spikelet meristem (upper ridge), X= repressed lower ridge, and FM= floret meristem. The numbers following the capital W represent different stages of wheat spike development using the Waddington scale: W1.0= vSAM (**d**, left), W1.5= initial transition of the vSAM to an IM (shown in Fig. 2). W2.0= early double ridge stage (**d**, center). W3.0= glume primordium (**e**). W3.5= lateral FMs subtended by lemma primordia and IM transition into a SM (IM→TS, **d**, right). (**f-h**) Wheat mutants with altered spike architecture. (**f**): *vrn1 vrn2*-null with increased SNS (**g**) *svp1 vrt2*-null mutants with increased SNS and faster development of basal spikelets. **(h)** Branched spike of “Miracle wheat”.

The spike architecture is determined by the changes in activity and identity of different meristems and by the timing of these changes. In wheat, both vegetative shoot apical meristems (vSAM) and IM produce lateral meristems in an alternate-distichous pattern, but these meristems differ in activity and identity. In the vSAM, bract meristems develop rapidly into leaves, whereas their axillary meristems (AxM) are temporarily repressed before initiating tillers. By contrast, in lateral meristems generated by the IM, the lower ridges (bract meristems) are repressed, while the upper ridges (equivalent to AxM) rapidly develop into spikelets (Fig. 1d). The developmental stages of the wheat spikes are described using the Waddington scale [5] (see Fig. 1 legend).

In wheat, spikelet development proceeds faster at the central part of the spike, resulting in the characteristic lanceolate shape of the wheat spike (Fig. 1a and e). By the time the central spikelets start forming floret meristems (FMs), the inflorescence meristem (IM) transitions into a spikelet meristem (SM) that forms a terminal spikelet (henceforth IM→TS) with a different orientation from the lateral spikelets (Fig. 1d). Several wheat mutants have been recently identified that affect the number of spikelets per spike (Fig. 1f), the shape of the spike (Fig. 1g), or the presence of supernumerary spikelets or branches (Fig. 1h).

This review summarizes the current knowledge of genes and gene networks that regulate wheat spike development, from the initial vSAM→ IM transition to the IM→TS transition (Fig. 1d), with a focus on the regulation of the spikelet number per spike (SNS). The first section describes the role of key MADS-box genes in the regulation of the early stages of spike development, whereas the second section covers genes that specifically affect SNS by regulating the rate at which SMs are produced or the timing of the IM→TS transition (Fig. 1d-g). The third section discusses an alternative way to increase SNS by generating supernumerary spikelets (SS) or branches (Fig. 1h). Finally, this review integrates recent advances in spatial transcriptomics and single-cell expression analyses and discusses the potential contributions of these new technologies to our understanding of wheat spike development and the improvement of wheat productivity.

Due to space limitations, the regulation of floret fertility and grain number per spikelet is not covered in this review, although its agronomic importance is recognized. Also excluded are the roles of phytohormones, nutrients, and stresses in spike development, which have been comprehensively reviewed elsewhere [6,7].

### 1. The critical role of MADS-box genes in the initial stages of wheat spike development

Wheat spike development starts with the vSAM→IM transition, which is driven by the upregulation of the *VERNALIZATION1* (*VRN1*) gene (Fig. 2a and b). However, *VRN1* upregulation in the vSAM is not sufficient to complete spike development and stem elongation, which requires long-day (LD) induction of *FLOWERING LOCUS T1* (*FT1*) in the leaves. The FT1 protein (florigen) is transported to the developing spike, where it simultaneously up-regulates *VRN1* and gibberellin (GA) biosynthetic genes [8] (Table S1). Both GA and *VRN1* contribute to the transcriptional up-regulation of *SUPPRESSOR OF OVEREXPRESSION OF CONSTANS1* (*SOC1*) and *LEAFY* (*LFY*) [8], two important regulators of inflorescence development. Another important role of VRN1 in the leaves is to repress the LD flowering repressor *VRN2* and the age pathway repressor *AP2L1*, both of which are negative regulators of *FT1* [9,10].

**Fig. 2.**
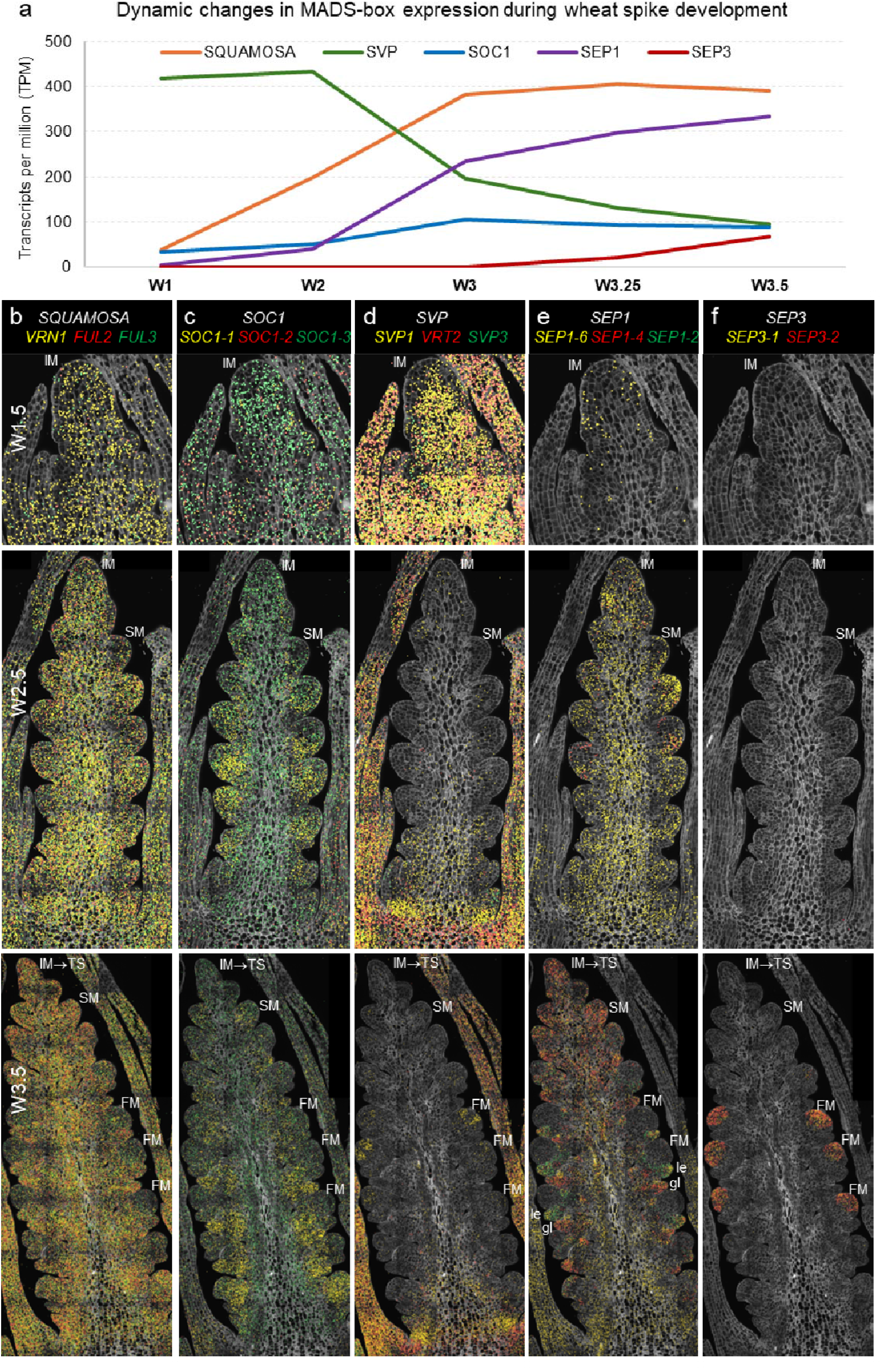
Dynamic changes in MADS-box gene expression during wheat early spike development. **(a)** Curves are based on RNA-seq data of wheat spike development in tetraploid wheat Kronos [18]. Individual points are the sum of TPM of all homeologs and paralogs within each MADS-box group (Table S1-2). **(b-f)** Spatial expression of MADS-box gene families across three spike developmental stages [19]. Cell walls are stained with calcofluor white, and dot colors correspond to the gene name colors in the same panel. IM= inflorescence meristem, SM= spikelet meristem, FM= floral meristem, le= lemma, gl= glume. **(b)** *<u>SQUAMOSA</u>. VRN1* is expressed earlier than the other two *SQUAMOSA* genes. All three genes are expressed at relatively lower levels at the FMs. **(c)** *<u>SOC1</u>. SOC1-2* expression is highest at W1.5 and then decreases, whereas *SOC1-1, SOC1-3*, and *SOC1-5* expression increase during early spike development (*SOC1-4* was not detected). **(d)** *<u>SVP</u>*. Genes are expressed in the IM at W1.5 and at the base of the spike at W2.5 and W3.5. *SVP1* is also expressed in developing anthers. **(e)** *<u>SEP1</u>. SEP1-6* is the earliest expressed gene in this clade, *SEP1-4* is enriched in the distal part of the spike and glume primordia, and *SEP1-2* is enriched in lemma primordia. **(f)** *<u>SEP3</u>*. *SEP3-1* and *SEP3-2* are expressed in FMs together with other floral homeotic genes (Table S1).

*VRN1* and its closest paralogues *FUL2* and *FUL3* are MADS-box genes of the *SQUAMOSA* clade that are widely expressed during early spike development (Fig. 2a-b). Spikes of plants carrying knockout mutations in all *VRN1* homeologs (*vrn1*-null) are normal [9], but *vrn1-*null combinations with *ful2-*null and *ful3-*null mutations (*SQUAMOSA-*null) result in abnormal spikes where spikelets are replaced by vegetative tillers subtended by leaves [4]. These *SQUAMOSA*-null plants (developed in a genetic background lacking the functional flowering repressor *VRN2* to avoid extremely late heading) still undergo a vSAM→IM transition followed by a double-ridge stage, elongated stems, and rapidly developing AxM in the “spike” region [4]. These results indicate that the *SQUAMOSA* genes play essential roles in suppressing the lower ridge and conferring spikelet meristem identity to the upper ridge, but that they are not essential for the initiation of the inflorescence scaffold [4].

The genes responsible for the initiation of inflorescence development in the wheat *SQUAMOSA*-null mutants are currently unknown, but based on evidence from Arabidopsis and rice, MADS-box genes from the *SOC1* and *SHORT VEGETATIVE PHASE* (*SVP*) clades are strong candidates. In Arabidopsis, genes from these two clades contribute to early inflorescence development by directly regulating *LFY* [11-13]. In rice, the *SOC1* homolog *OsMADS50* (ortholog of wheat *SOC1-3*) promotes the transition to the reproductive phase [14], suggesting a conserved function.

Among the five *SOC1* paralogs identified in wheat [15], the expression of *SOC1-1, SOC1-3*, and *SOC1-5* are significantly upregulated during the vSAM to IM transition (Table S2 and Fig. 2a), which is consistent with a role in early inflorescence development. In addition, *SOC1-1* expression is upregulated by the addition of GA, which accelerates spike development [8]; and the SOC1-3 protein competes with VEGETATIVE TO REPRODUCTIVE TRANSITION 2 (VRT2, SVP clade) for interaction with VRN1 to regulate heading time [16]. The wheat *SOC1* paralogs exhibit different spatial expression profiles during wheat spike development (Fig. 2c), but the elucidation of their specific functions will require further experimental validation.

The wheat SVP transcription factors, which include SVP1, VRT2, and SVP3, physically interact with the SQUAMOSA proteins [17]. Combined loss-of-function mutations in *SVP1* and *VRT2* genes delay heading time, increase spikelet number per spike (SNS, Fig. 1f-g), and reduce plant height in a similar way to the *vrn1-*null mutants. The similar effects of these mutants together with the physical interactions between their encoded proteins suggest that SQUAMOSA and SVP work cooperatively to regulate early spike development [17].

The wheat *SVP* genes are co-expressed with the *SQUAMOSA* genes in the early IM at W1.5, but at later stages their expression becomes restricted to the base of the spike and to the leaves (Fig. 2d). The transcriptional downregulation of the *SVP* genes by the *SQUAMOSA* genes is important for the normal progression of spike development, as SVP proteins can interfere with the SQUAMOSA-SEPALLATA1 (SEP1) protein interactions [17]. Plants carrying high-expressing natural *VRT2* alleles or transgenic plants overexpressing this gene exhibit more vegetative traits such as larger glumes and lemmas [17,20]. By contrast, the vegetative characteristics of the *vrn1 ful2* spikes are mitigated when combined with *vrt2* mutations [17], highlighting the importance of the downregulation of the *SVP* genes for normal spike development.

The *SEP1* and *SVP* genes show opposite expression profiles (Fig. 2a). Among the *SEP1* genes, *SEP1-6* expression is initiated the earliest (Fig. 2e), and its overexpression in wheat results in reduced SNS [21]. Combined mutations in the rice homologs of *SEP1-6* (*OsMADS34*) and *SEP1-4* (*OsMADS5*) lead to enhanced branching in rice panicles [22], but have limited effects on barley spikes [23,24]. Mutations in the *SEP1-2* ortholog in rice (*OsMADS1*) result in floral organs with leaf-like characteristics [25], whereas mutations in the barley ortholog lead to smaller lemmas and awns at room temperature [26] and branched spikes at high temperatures [23]. Given the different effects of the *SEP1* mutants in rice and barley, it is not possible to infer their function in wheat without dedicated research efforts. The *SEP3* genes are co-expressed in the FM (Fig. 2f) with other floral homeotic genes [19] and contribute to floral organ identity [27].

MADS-box proteins form quaternary complexes of varied compositions, each with specific DNA binding specificities [28]. Therefore, the early upregulation of *VRN1* in the IM of wheat (Fig. 2b) and the gradual changes in the relative expression of other MADS-box genes (Fig. 2a) likely affect the composition of these quaternary complexes and the sequential morphological changes observed during the early stages of wheat spike development.

### 2 Regulation of wheat spike determinacy and spikelet number per spike (SNS)

#### 2.1 Regulation of wheat spike determinacy

Both wheat and barley produce spikes with a genetically regulated number of spikelets, but they differ in the fate of the IM. In the determinate wheat spike, the IM transitions into an SM and forms a terminal spikelet (Fig. 1a), whereas in the indeterminate barley spike, the IM is exhausted and eventually dies. However, there are mutation that can revert these phenotypes. In barley, mutations in *APETALA2-LIKE5* (*ap2l5*-null) result in a determinate spike [29], whereas in wheat, combined loss-of-function mutations in *VRN1* and *FUL2* result in an indeterminate inflorescence [4].

In wheat, loss-of-function mutations in *AP2L5* lead to reduced SNS [30], whereas increased *AP2L5* expression increases SNS [31]. However, neither of these changes affects wheat spike determinacy. In both species, mutations in *AP2L5* result in an increased number of florets per spikelet, suggesting similar functions in spikelet development but distinct roles in spike determinacy.

#### 2.2 Regulation of spikelet number per spike

Individual loss-of-function mutations in *VRN1* or *FUL2* significantly increase SNS, but the effects are more pronounced in *vrn1-*null (58%) than in *ful2*-null mutants (10%) [4]. Both *vrn1-* null (Fig. 1f) and *svp1 vrt2-*null (Fig. 1g) mutants show delayed spike and spikelet development associated with a reduced rate of SM production and a significantly delayed IM→TS transition (Fig. 3a-b). In Arabidopsis, MADS-box tetramers including SQUAMOSA proteins act as pioneer transcription factors that bind closed chromatin and modify it to an open state, ensuring access to other transcription factors that specify cell or organ fate [32]. We hypothesize that this may explain how the brief co-expression of SQUAMOSA and SVP proteins in the early IM (W1.5, Fig. 2b and d) is sufficient for a persistent change in the rate of SM production (Fig. 3a-b).

**Fig. 3.**
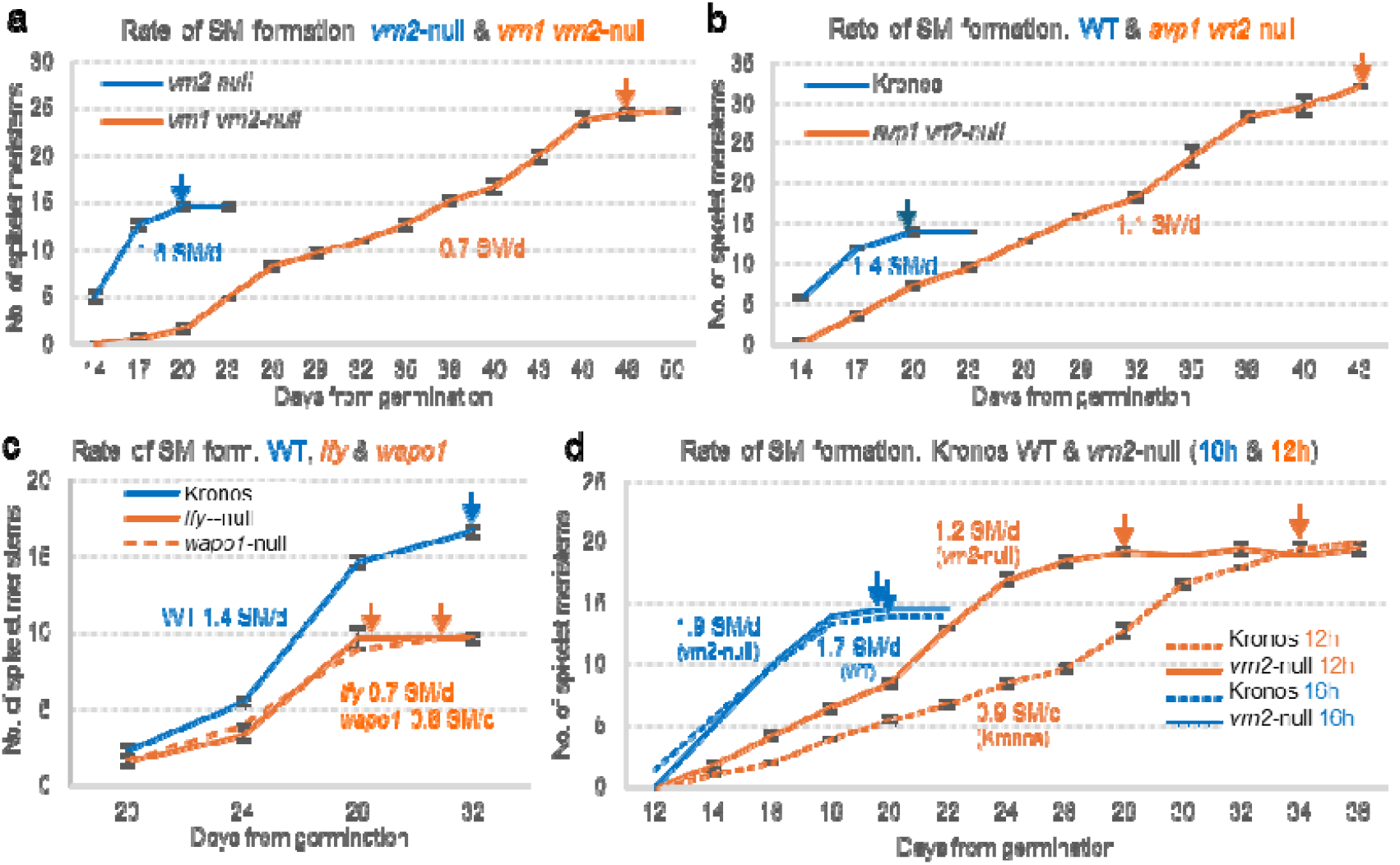
Rate of spikelet meristem (SM) formation and IM→TS transition across mutant backgrounds and photoperiods. **(a)** *vrn1 vrn2-*null mutant versus *vrn2-*null control. **(b)** *svp1 vrt2-*null mutant versus Kronos control. **(c)** *lfy-*null and *wapo1-*null mutants versus Kronos control [33]. **(d)** Effect of photoperiod on SM production rate: 16h light /8 h darkness (blue) vs. 12h light / 12h darkness (orange) in Kronos (dotted lines) and Kronos *vrn2-*null mutant (solid line). Longer photoperiods [40] and absence of *vrn2* [9] both result in increased transcript levels of *FT1* and a faster rate of SM production. Arrows indicate the approximate timing of the IM→TS transition. Rates are expressed as the number of SMs formed per day. Data points represent the averages of three dissected apices and error bars indicate standard errors of the means (SEM). Data is available on Table S3.

The rate of SM production is also reduced relative to the wildtype in the *lfy* and *wapo1* mutants, but the timing of the IM→TS transition is not significantly altered [33] (Fig. 3c). Loss-of-function mutations in *LFY, WAPO1*, or both result in similar reductions in SNS, suggesting that they work cooperatively to regulate this trait. This hypothesis is supported by the physical interaction between their encoded proteins and their co-expression in the early IM at W1.5 [33]. *LFY* and *WAPO1* are not co-expressed in the IM at later stages of spike development, but since *LFY* is a pioneer transcription factor that can bind closed chromatin and promote an open state [34], this brief interactions may be sufficient to trigger the persistent effects on the IM rate of SM production observed in Fig. 3c [33]. Natural alleles with positive effects on SNS have been identified for *WAPO1* [35] and *LFY* [36] and both show evidence of positive selection during wheat breeding history and a significant genetic interaction for SNS.

The IM→TS transition at W3.5 is characterized by the upregulation of *SQUAMOSA* genes *VRN1* and *FUL2* and the downregulation of both *LFY* [33] and the *SQUAMOSA PROMOTER BINDING PROTEIN-LIKE* gene *SPL14* in the IM region [19]. Loss-of-function mutations in *SPL14* result in a premature IM→TS transition and a 42% reduction in SNS, demonstrating an important role of this gene in spike development [19]. This hypothesis is further supported by the discovery of natural alleles for *SPL14* and its close paralog *SPL17* with positive effects on wheat SNS [37,38]. An RNA-seq study of early spike development comparing wildtype wheat Fielder with a combined *spl14 spl17-*knockout mutant, suggests that *SPL14* and *SPL17* act as promoters of *SOC1-1* and *SOC1-5* and repressors of *SQUAMOSA* and *SVP* MADS-box genes [37].

Another group of genes with significant effects on wheat SNS includes the *PHOTOPERIOD1* (*PPD1*) gene and its downstream targets *FT1* and *FT2*. Plants carrying the *PPD1* photoperiod*-* sensitive (PS) allele have lower *FT1* expression and higher SNS than those with photoperiod*-* insensitive (PI) alleles. Consistently, wheat plants with loss-of-function mutations in *PPD1* show reduced *FT1* expression and higher SNS [39,40]. Shorter days also reduce *FT1* transcript levels, delay spike and spikelet development, and increase SNS (Fig. 3d) [40]. These effects are attenuated in the *vrn2-*null mutants, which exhibit higher *FT1* expression and a faster rate of SM formation (Fig. 3d). The dominant role of florigen proteins in the regulation of SNS is also supported by the low SNS observed in plants overexpressing FT1 [41], FT2 [42], or FT3 [43]. Based on these results, we hypothesize that the delayed IM→TS transition in the *vrn1 vrn2-*null and *svp1 vrt2-*null mutants (Fig. 3a-b) could be associated with the downregulation of *FT1* observed in the leaves of these mutants [4,17].

The *FT2* gene also affects SNS, and its encoded protein physically interacts with bZIPC1, a homeodomain-leucine zipper transcription factor [40,44]. The two genes affect SNS in opposite directions, with loss-of-function mutations in *FT2* increasing SNS and those in *bZIPC1* reducing SNS. Natural alleles with positive effects on SNS and evidence of positive selection during wheat improvement have been identified for both genes [44,45]. Both *FT2-bZIPC1* and *WAPO1-LFY* showed significant genetic interactions for SNS that have led to the identification of optimal allele pairs that maximize SNS [36,45]. These results illustrate the practical value of understanding the gene networks and epistatic interactions that regulate spike development.

A wheat spike transcriptome association study identified additional transcription factors that influence SNS and validated them using transgenic approaches [21]. Over-expression of *SEP1-6* (synonymous *PAP2*) or *GRAIN NUMBER INCREASE 1* gene (*GNI1*, ortholog of barley *VRS1*) reduced SNS in a dosage-dependent manner, whereas overexpression of *CENTRODIALIS 2* (*CEN2*, synonymous *TFL-D1*) increased SNS [21]. An important challenge in the coming years will be to elucidate the interactions among the different genes affecting SNS.

Finally, the number of fertile spikelets can be also affected by the proportion of basal rudimentary spikelets. High *VRT2* levels are associated with a higher number of basal rudimentary spikelets, suggesting that *VRT2* can delay the development of basal spikelets and contribute to the lanceolate shape of the wheat spikes [46]. This hypothesis is supported by the triangular spike shape of the *svp1 vrt2* mutant at W3.0 (Fig. 1g), where basal spikelet development is not delayed relative to central spikelets, unlike in the wildtype (Fig. 1d).

Increases in SNS do not guarantee increases in grain number per spike (GNS) because floret degeneration can offset the gains associated with additional spikelets. Fortunately, a natural allele of *GNI1* provides a tool to increase spikelet fertility and reduce premature floret abortion [47]. Even when increases in SNS are translated into more GNS, reduced grain weight can offset the positive effect of increased GNS on grain yield. When plants cannot produce sufficient resources to fill the extra grains, a negative correlation is usually observed between GNS and grain weight. For example, the favorable *WAPO1* allele for increased SNS showed positive effects on grain yield only when introgressed into high-biomass genotypes and when lines were grown under well-fertilized and well-watered conditions [35,48]. These experiments suggest that balanced increases in both sink and source, together with favorable environmental conditions, are necessary to translate increases in SNS into higher grain yields.

### 3. Regulation of branching in the wheat spike

An alternative strategy to improve wheat SNS is to increase the number of spikelets produced per node, generating supernumerary spikelets (SS). In mutants such as ‘Miracle-Wheat’ (Fig. 1h) and ‘Compositum-Barley’ (*com2*), basal spikelets can be replaced by branch-like structures resembling smaller secondary spikes (coflorescences) [49,50], suggesting that these lateral meristems acquired an IM identity before transitioning to a terminal spikelet.

Branched-like phenotypes and SS have been associated with hypomorphic mutations in the APETALA2/ETHYLENE RESPONSE FACTOR (AP2/ERF) domain transcription factor FRIZZY PANICLE (FZP) [49-51]. However, in more severe *FZP* mutants in rice [52] and durum wheat [19], secondary axes emerge from the axils of the glumes, forming new bracts/glumes that produce tertiary axes and repeat the process. These recurrent glume structures suggest that, in addition to its role in suppressing branching, an important function of *FZP* is to suppress the glume’s AxM. This result is consistent with the restricted expression of *FZP* in the axils of wheat glume primordia (Fig. 4a-b).

**Fig. 4.**
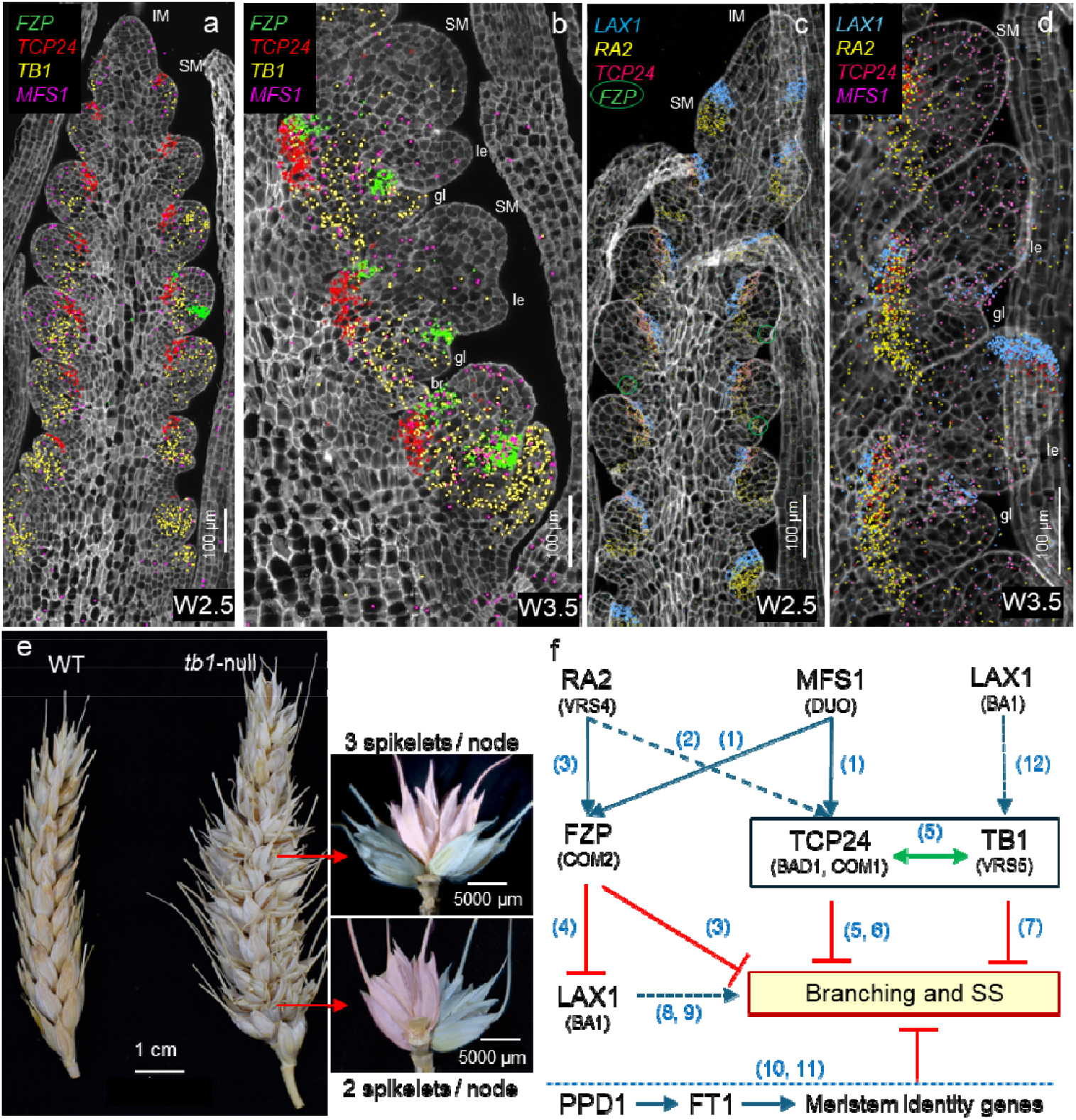
Genes and gene network regulating branching. (**a-d**) Single-molecule fluorescence *in situ* hybridization of Kronos developing spikes at W2.5 (**a** and **c**) and W3.5 (**b** and **d**). Cell walls are stained with calcofluor white, and dot colors correspond to gene name colors in the same panel. gl= glume, le= lemma, br= suppressed bract, SM= spikelet meristem, IM= inflorescence meristem. (**a-b**) Expression profiles of *FZP, TCP24, TB1* and *MFS1* based on [19] (**c-d**) Expression profiles of *LAX1, RA2, TCP24* and *MFS1* in Kronos spikes based on a 10X-Xenium spatial transcriptomics study. (**c**) Incipient expression of *FZP* is indicated by green circles (**e**) Left: Spikes of wildtype Kronos and *tb1-*null CRISPR mutant showing SS. Right: nodes bearing two or three spikelets. Note the different orientation of the terminal (pink) and lateral spikelets (blue), their similar development, and the higher frequency of triplets at the center of the spike. (**f**) Working model of a gene network regulating branching in wheat. Blue arrows indicate promotion, red T-shaped symbols indicate repression, and green double arrows represent protein-protein interactions. Solid lines are supported by wheat or barley studies and dotted lines by studies in the more distantly related rice or maize. Gene names and IDs are included in Table S1. Numbers in parenthesis indicate references: 1:[53], 2:[55], 3:[50], 4:[51], 5:[56], 6:[57], 7:[58], 8:[59], 9:[60], 10:[61], 11:[62], 12:[60].

Studies in wheat and barley suggest that *FZP* transcript levels are positively regulated by two transcription factors: wheat MULTI-FLORET SPIKELET1 (TaMFS1, also referred to as DUO [53]) and barley RAMOSA2 (HvRA2, synonymous VRS4) [50]. In wheat, MFS1 directly promotes the expression of *FZP* [53] and the expression domains of these two genes show significant overlap at early stages of spikelet development (Fig. 4b). Consistent with its effect on *FZP*, the wheat *mfs-B1* mutant also produces spikes with SS [53].

In barley, loss-of-function mutations in *HvRA2* (*VRS4*) are also associated with reduced *FZP* transcript levels. These mutants also promote lateral spikelets, occasional branch-like formation [50], and reduced expression of barley *VRS1* and *SISTER OF RAMOSA3* [54]. In the wheat spike, *RA2* is first induced in the abaxial part of initiating SMs, then surrounds the growing SM (Fig. 4c), and later delimits the base of the developing spikelet, overlapping with the *FZP* expression domain in the axil of the second glume (Fig. 4d).

Wheat MFS1 [53] and maize RA2 [55] are also positive regulators of *TCP24*, which corresponds to *COMPOSITUM1* (*COM1*) in barley and *BRANCH ANGLE DEFECTIVE 1* (*BAD1*) in maize [55]. Loss-of-function mutations in *COM1* result in branched barley spikes, suggesting that the active COM1 promotes SM identity and contributes to coflorescences suppression [56,57]. The interactions among *MFS1, RA2*, and *TCP24* in wheat are also supported by significant correlations among their expression profiles in a single-cell RNA-seq (scRNA-seq) study of early wheat spike development [19] and by their overlapping expression domains (Fig. 4d). Moreover, increased expression of all three *TCP24* homeologs is observed in the wheat *mfs-B1* mutant [53]. A similar interaction has been observed in maize, where *RA2* operates upstream of *BAD1* [55].

TCP24 belongs to the CYC/TB1 clade of TCP transcription factors, which also includes TCP22, TEOSINTE BRANCHED1 (TB1), and TB2. In barley, TB1 is known as INTERMEDIUM SPIKE-C (INT-C or VRS5), and the mutant allele contributes to the six-rowed phenotype [63]. In wheat, *TB1* expression is detected in the SMs at W2.5 and becomes restricted to the base of the spikelets at W3.5, and at both stages is more abundant at the base of the spike (Fig. 4a-b). Knockout mutations of both *TB1* homeologs in tetraploid wheat (*tb1-*null) result in frequent SS (Fig. 4e, Table S4), as reported previously [58]. These results suggest that *TB1/INT-C* modulates SM identity, thereby contributing to the repression of SS in both barley and wheat. This result is consistent with the role of *TB1* in maize, where it acts as a repressor of axillary branching [64]. Surprisingly, *TB1* duplications or transgenic overexpression were also associated with SS [62], suggesting that balanced levels of *TB1* may be required to prevent SS formation.

In maize, *TB1* transcription is regulated by the bHLH transcription factor *BARREN STALK1* (*BA1*), the ortholog of rice *LAX PANICLE1* (*LAX1*) [60]. The *LAX1* ortholog in wheat is expressed at the adaxial boundary layer of the SMs from the earliest stages of SM formation (Fig. 4c). A similar adaxial position has been reported in rice [59] and maize [60], suggesting a conserved function. *BA1* acts downstream of auxin signaling to position boundary regions for AxM formation [65]. Loss-of-function mutations in *BA1* or *LAX1* result in drastic reductions in panicle branching and lateral spikelet formation in both maize and rice [59,60], whereas mutations in the wheat orthologs produce milder effects [66]. Wheat plants with mutations in all three *LAX1* homeologs have a more compact spike [66] that resembles gain-of-function mutations in the domestication gene *Q* (*AP2L5*) [31,67]. By contrast, overexpression of wheat *LAX1* reduces spike threshability and spikelet density, like the pre-domestication *q* allele [66]. LAX1 and Q proteins physically interact with each other, providing a potential mechanism for their opposite effects on these domestication traits [66]. In wheat, FZP has been reported to directly repress *LAX1* transcription by binding to its regulatory regions [51]. However, the functional significance of this interaction remains unclear, given that *LAX1* is highly expressed from the very early stages of SM development, whereas *FZP* is detected only in the more mature SMs at W2.5 (Fig. 4c). *FZP* expression becomes stronger at later stages (W3.0 and W3.5), when glume primordia are well developed (Fig. 4b). A working model summarizing the interactions among the different genes regulating SS is presented in Fig. 4f.

In hexaploid wheat, a particular class of SS, designated as paired spikelets (PS), is characterized by the formation of a secondary spikelet immediately adjacent to and below (abaxial) a more advanced main spikelet. At maturity, PS appear parallel to each other and are more frequent at the central nodes of the spike. PS are affected by genes in the photoperiod pathway, with shorter photoperiods, mutations in *PPD1*, and reduced expression of *FT1* all increasing PS frequency [61]. Since these changes are associated with reduced expression of meristem identity genes (*VRN1, FUL2*, and *FUL3*), the authors of [61] hypothesized that this reduction may contribute to a brief reversion to IM identity or a weaker SM identity, resulting in PS formation.

In summary, multiple mutants generate SS or coflorescences, likely as a result of modifications in SM identity. Considering that spikelets are small inflorescences, transitions between IM and SM in either direction (IM→SM or SM→IM) likely require less gene regulatory changes than the vSAM→IM transition or the initiation of FMs. We hypothesize that a short or incomplete SM→IM transition could generate PS with a dominant terminal spikelet and a secondary lateral spikelet, whereas a longer and/or more complete SM→IM transition could allow for the formation of one or more fully developed lateral spikelets. An even longer SM→IM transition could generate coflorescences as observed in Miracle Wheat. It remains unclear why some mutations induce higher SS frequencies at the center of the spike (e.g. *TB1*), whereas others result in coflorescences at the base of the spike (e.g. *FZP*). Studying branching in the un-branched *Triticeae* spikes can provide valuable insights into grass inflorescence development and evolution.

## Conclusions and perspectives

Increasing spikelet number through SS or branching has been an attractive strategy to increase grain yield, but initial efforts using the *FZP* natural mutants have been limited by trade-offs in fertility and grain size. A recent study using a cross between a spike-branching landrace and the elite durum cultivar CIRNO showed that breeding efforts to generate a better balance of source– sink traits can reduce yield penalties associated with branched phenotype [68]. However, developing higher yielding wheat varieties with SS or branched spikes will require a dedicated breeding effort to fine-tune the multiple epistatic interactions that exist within the complex gene network regulating branching (Fig. 4f). The *GNI1* allele for high fertility [47] may provide an alternative strategy to mitigate the reductions in SS fertility, whereas high-biomass varieties with higher source resources may reduce the effect of negative correlations between grain number and grain size.

Meanwhile, rapid progress has been made in the identification of genes that control grain number by modifying the rate of SM production, the timing of the IM →TS transition, and/or the number of fertile florets. Favorable alleles and allele combinations are being identified and incorporated into breeding programs through marker-assisted selection or by integrating functional SNPs into high-throughput marker platforms used for genomic selection. The impressive number of spikelets and grains produced by high yielding triticale varieties suggests that there is still room for improvement of the SNS trait in wheat.

The identification and functional characterization of genes regulating wheat spike development have been greatly accelerated in recent years by the development of new tools including CRISPR, sequenced mutant populations, improved wheat transformation efficiency, and sequenced genomes. These resources have been complemented by single-cell RNA-seq and spatial transcriptomic studies, which provide the cellular resolution required to visualize the rapid changes in gene expression that occur during spike development [19,69]. These studies can accelerate the discovery of new genes affecting spike architecture by identifying genes that are co-expressed in the same cells or share similar spatial and temporal expression profiles with genes of known functions [19]. Haplotype analysis of the new genes and the association of these haplotypes with available wheat GWAS and QTL studies has the potential to identify valuable natural variants for wheat improvement.

In parallel, multi-omics studies are integrating expression profiles, chromatin accessibility, binding motifs, GWAS, and evolutionary information into comprehensive networks that will help prioritize the functional characterization of novel regulators of wheat spike development [70,71]. The knowledge generated by these new tools can enhance our ability to engineer more productive wheat spikes.

Each year, more than 200 trillion wheat spikes are harvested globally, providing one fifth of the calories and proteins consumed by the human population. Therefore, every small improvement in wheat spike productivity is worth the effort.

## Acknowledgements

We are grateful to Dr. Thorsen Schnurbusch (Leibniz Institute of Plant Genetics and Crop Plant Research, IPK) for the image of Miracle-Wheat (Fig. 1h). We also appreciate the insightful comments and suggestions from Dr. Thorsen Schnurbusch and Dr. Scott Boden (The University of Adelaide) on the branching section of this review, and from Dr. Francois Parcy (Universite Grenoble Alpes) on comparative inflorescence development.

## Funding statement

This project was supported by the Agriculture and Food Research Initiative Competitive Grant 2022-68013-36439 (WheatCAP) from the USDA National Institute of Food and Agriculture, and by the Howard Hughes Medical Institute.

References and recommended reading Papers of particular interest, published within the period of review, have been highlighted as: * of special interest and ** of outstanding interest

## Notes

### Competing Interest Statement

The authors have declared no competing interest.

### Summary of Updates

This manuscript has been revised to incorporate reviewers comments. We incorporated a new Fig. 1d describing meristem transitions during spikelet development. We also proposed a mechanism to explain how interacting transcription factors that are co-expressed only at the early stages of spike development (W1.5) can have a more persistent effect on the rates of spike development. We introduce the term coflorescences to designate branches that resemble the main inflorescence in Miracle Wheat. We presented a hypothesis for the origin of different types of supernumerary spikes. We added examples on how the multi-omics data can be used to identify allelic variation useful for wheat improvement.

